# Robust Design for Coalescent Model Inference

**DOI:** 10.1101/317438

**Authors:** Kris V Parag, Oliver G Pybus

## Abstract

—The coalescent process describes how changes in the size of a population influence the genealogical patterns of sequences sampled from that population. The estimation of population size changes from genealogies that are reconstructed from these sequence samples, is an important problem in many biological fields. Often, population size is characterised by a piecewise-constant function, with each piece serving as a population size parameter to be estimated. Estimation quality depends on both the statistical coalescent inference method employed, and on the experimental protocol, which controls variables such as the sampling of sequences through time and space, or the transformation of model parameters. While there is an extensive literature devoted to coalescent inference methodology, there is surprisingly little work on experimental design. The research that does exist is largely simulation based, precluding the development of provable or general design theorems. We examine three key design problems: temporal sampling of sequences under the skyline demographic coalescent model, spatio-temporal sampling for the structured coalescent model, and time discretisation for sequentially Markovian coalescent models. In all cases we prove that (i) working in the logarithm of the parameters to be inferred (e.g. population size), and (ii) distributing informative coalescent events uniformly among these log-parameters, is uniquely robust. ‘Robust’ means that the total and maximum uncertainty of our estimates are minimised, and are also insensitive to their unknown (true) parameter values. Given its persistence among models, this formally derived two-point theorem may form the basis of an experimental design paradigm for coalescent inference.

---

The coalescent process [1] is a popular population genetic model that describes how past changes in the size or structure of a population shape the reconstructed (observed) genealogy of genetic sequences sampled from that population. This genealogy is also known as a coalescent tree or phylogeny. The estimation of a function that describes past population size from the sequences, or from a reconstructed phylogeny, is a problem encountered in many fields including epidemiology, conservation and anthropology. Accordingly, there is an extensive and growing literature [2] [3] [4] [5] [6] [7] [8] [9] [10] [11] [12] [13] focussed on developing new statistical methods for solving coalescent inference problems.

However, the power and accuracy of the resulting coalescent estimates is not solely a function of the statistical method employed. Variables under the control of the experimenter, such as choices of where and when sequences are sampled, or on how time is discretised, may have a strong influence on the performance and reliability of coalescent inference methods [14] [9] [11]. Good designs can result in sharper inferences and sounder conclusions [14], whereas bad designs, such as size-biased sampling strategies, can lead to overconfident or spurious estimates [15]. The best approach to coalescent inference should therefore jointly optimise experimental design and statistical methodology.

Surprisingly few studies have investigated optimal coalescent inference design. These works [14] [16] [12] [17] [15], typically take a simulation based approach, in which several alternate designs are numerically examined and compared. While such studies can yield useful hypotheses about the components of good designs, they can neither provide analytic insights nor criteria for provably optimal experimental design. A more general and methodical analysis is therefore warranted.

Additionally, there has been little consideration of what data or parameter transformations might aid experimental design. This contrasts with the development of inference theory in other fields. For example, in regression problems, research has emphasised the benefits of power transformations and regularisation procedures [18]. While some coalescent inference methods have used parameter transformations (e.g. the log transform), these are usually justified for heuristic or practical reasons, such as algorithmic stability or ease of visualisation [12] [19]. As a result, parameter transformations used in the coalescent literature are applied inconsistently and rigorous proof of their benefits is lacking.

Here we take a fully analytical approach and formally derive optimal design criteria for coalescent inference. As we are interested in widely applicable theoretical insights, we do not construct method-specific rules, but instead define benchmarks which, if achieved, guarantee certain well-defined and desired properties. We investigate three popular coalescent models, which we class as ‘piecewise’ due to the characteristic functions they infer. For each model we describe a coalescent tree as being composed of sample lineages, with time flowing from the present into the past. A coalescent event occurs when two lineages merge into a single ancestral lineage.

(1) Skyline demographic models. These infer past population size changes using piecewise-constant, time-varying functions [20], and usually feature genealogies with many samples from a few (usually one) loci [13]. The large sample size of these trees means that the choice of sequence sampling times is a critical design variable that can significantly influence the precision of population size inference. Skyline models are popular in epidemiology where the population describes the number of infected individuals in an epidemic. Optimal sampling designs could improve epidemic surveillance and control strategies [4] [14].

(2) Structured coalescent models. Here the population is divided into distinct but connected sub-populations (demes), which typically represent different spatial locations. Usually each deme has a constant (stable) population size. Lineages may migrate between demes but can only coalesce within demes. The population sizes and migration rates are our parameters of interest [21] [22]. The locations and times of sampled sequences, which are our design variables here, are known to bias inference under these models [9]. This model has been applied to describe the migration history of animal, plant and pathogen populations [9].

(3) Sequentially Markovian coalescent (SMC) models. These are typically applied to complete metazoan genomes, and consider many independent coalescent trees (multiple unlinked loci), each containing few (or two) samples. SMC processes involve recombination, and event times are discretised to occur in finite intervals. Past population size change is often assumed to be piecewise-constant and most SMC applications focus on human demographic history [10] [12]. The design variable here is the time discretisation, which controls the resolution with which population size changes are estimated. Poor discretisations can lead to overestimation or runaway behaviour [11].

We examine the above models using optimal design theory, which aims to optimise experimental protocols using statistical criteria that confer useful properties, such as minimum bias or maximum precision [23]. As the coalescent event times contain information about population size change, the distribution and total number of coalescent events controls the amount of information available. Within this context, we treat our sampling/discretisation choice problem as an experimental design on this coalescent event distribution.

We show that it is optimal to (i) estimate the logarithm of our parameters of interest, which usually refer to effective population size, and (ii) sample (through time and location) or discretise time such that coalescent events are divided evenly among each log scaled parameter. If (i)-(ii) are achieved, then the resulting design is provably robust, and optimal for use with existing maximum likelihood and Bayesian coalescent inference methods. ‘Robust’ means that both the maximum dimension and the total volume of the confidence ellipsoid that circumscribes (asymptotic) estimate uncertainty are jointly minimised. These two objectives hold for all piecewise coalescent models (such as those above) and therefore comprise simple, unifying rules for coalescent inference design.

In the Preliminaries we provide mathematical background on optimal design. We use these concepts to derive a robust design theorem for coalescent inference, in Results. This is then applied to each of the three coalescent models described above, yielding new and specific insights. We close with a Discussion of how our formally derived design principles relate to existing heuristics in the literature.

## Preliminaries

Consider an arbitrary parameter vector *ψ* = [*ψ*_1_,…, *ψ_p_*], which is to be estimated from a statistical model. Let 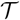 represent data (a random variable sequence) generated under this statistical model (the genealogy in the case of coalescent inference) and let 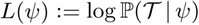 be the log-likelihood of 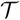 given *ψ*. The *p* × *p* Fisher information matrix, denoted 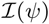, measures how informative 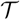 is about *ψ* [24]. Since all the coalescent models used here belong to an exponential family [25] (and so satisfy necessary regularity conditions [26]) then the (*i, j*)^th^ element of 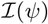 is 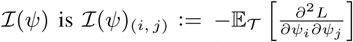, with the expectation taken across the data (tree branches). The Fisher information is sensitive to parametrisation choices. Eq. (1) gives the transformation between *ψ* and an arbitrary alternate parametrisation *σ* = [*h*(*ψ*_1_),…, *h*(*ψ_p_*)] = [*σ*_1_,…, *σ_p_*]. Here *h* is a differentiable function, with inverse *f* = inv[*h*] [25].

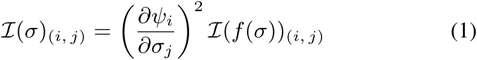

The Fisher information lower bounds the best unbiased estimate precision attainable, and quantifies the confidence bounds on maximum likelihood estimates (MLEs). For exponential families, these bounds are attained so that if 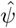 is the MLE then 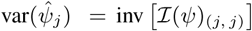 is the minimum achievable variance around the MLE of the *j*^th^ parameter by any inference method [27]. Importantly, for any given parametrisation, the Fisher information serves as a metric with which we can compare and rank various estimation schemes (e.g. different sampling or discretisation protocols).

Since all of our statistical models are finite dimensional, the Bernstein-von Mises theorem [28] [29] is valid. This states that, asymptotically, any Bayesian estimate will have a posterior distribution that matches that of the MLE, with equivalent confidence intervals, for any ‘sensibly defined’ prior. Such a prior has some positive probability mass in an interval around the true parameter value. As a result, Bayesian credible intervals also depend on the Fisher information and our designs are applicable to both maximum likelihood and Bayesian approaches to coalescent inference.

We now construct our piecewise coalescent experimental design problem. If the observed data 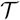 consists of *n* − 1 coalescent events (i.e. a tree with *n* tips) then the set {*m_j_*} for 1 ≤ *j* ≤ *p* with 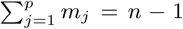 describes a coalescent event distribution. Here *m_j_* counts the coalescent events that are informative of parameter *ψ_j_*. Optimal designs are {*m_j_*} sets that satisfy desirable statistical criteria. This is illustrated for a two parameter skyline demographic model in Fig. 1 (with parameters *ψ_j_* = *N_j_*), for which sampling choices would be used to achieve the optimal {*m_j_*} sets. Statistical design criteria are typically functions of 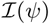, which defines our asymptotic uncertainty about 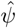. Geometrically, this uncertainty can be represented as a confidence ellipsoid centred on 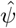 [30].

Designing the Fisher information matrix is equivalent to controlling the shape and size of this ellipsoid. We focus on two popular criteria, known as D and E-optimality [30] [23], the definitions of which are given in Eq. (2) and Eq. (3), with 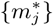 as the resulting optimal design. As we have *p* design variables (the *m_j_*), our confidence ellipsoid is *p*-dimensional. D-optimal designs minimise the volume of this confidence ellipsoid while E-optimal ones minimise its maximum diameter. Fig. 2 shows these ellipses for the skyline design problem of Fig. 1.

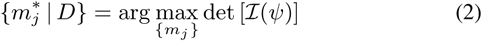

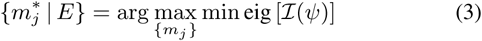

Here arg, det and eig are short for argument, determinant, and eigenvalues respectively. D-optimal designs therefore maximise the total available information gained from the set of parameters while E-optimal ones ensure that the worst parameter estimate is as good as possible [30] [23].

**Fig. 1:**
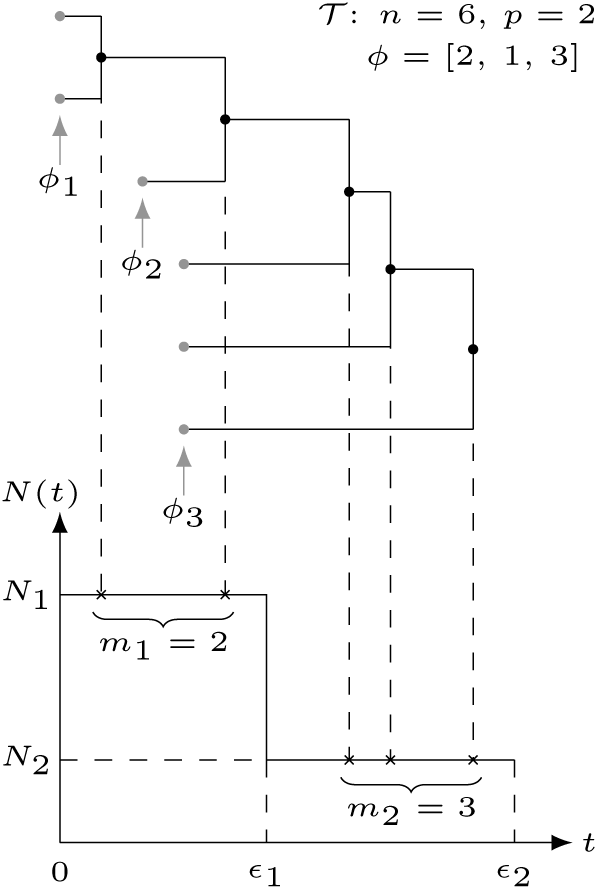
Two-parameter piecewise coalescent design problem. We show a *p* = 2 design problem for a skyline demographic coalescent model with population size parameters *N*_1_ and *N*_2_. An *n* =6 tip coalescent phylogeny, 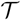, is shown with the *ϕ_k_* counting the samples introduced at the *k*^th^ sample time. The *j*^th^ population size parameter, *N_j_*, is only informed by the number of coalescent events, *m_j_*, occurring within its duration [*ϵ_j_*_−1_, *ϵ_j_*] (with *ϵ*_0_ = 0). We manipulate *ϕ* to achieve *m*_1_ and *m*_2_ counts that guarantee desirable properties for estimates of *N*_1_ and *N*_2_.

The above optimisation problems can be solved using majorization theory, which provides a way of naturally ordering vectors [31]. For some *p*-dimensional vectors 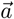 and 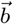, sorted in descending order to form 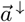 and 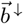, 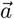 is said to majorize or dominate 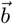 if for all *k* ∈ {1, 2,… *p*}, 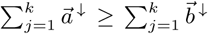 and 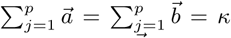. Here *κ* is a constant and this definition is written as 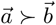 for short. The total sum equality on the elements of the vectors is called an isoperimetric constraint. Conceptually, 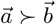 means that the elements of 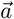 have the same mean as those of 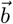, but a higher variance.

We will make use of Schur concave functions. A function *g* that takes a *p*-dimensional input and produces a scalar output is called Schur concave if 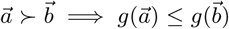. Importantly, it is known that the *p*-element uniform vector 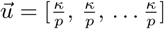 is majorized by any arbitrary vector of sum *κ* and dimension *p* [31]. This means that every 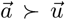. As a result, 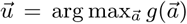 for any Schur concave function *g*. Thus if we can find a Schur concave function, and an isoperimetric constraint holds, then a uniform vector will maximise that function. This type of argument will underpin many of the following results.

## Results

### Naive Coalescent Design

We define a naive coalescent inference design as one that works directly in the original parametrisation of the model, which is usually effective population size or its inverse. Let *N* = [*N*_1_,…, *N_p_*] be the parameter vector to be estimated from a reconstructed genealogy, 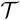. Defining 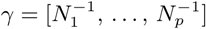, we will find that all three of the coalescent models we consider here have log-likelihoods, 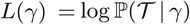, of the form of Eq. (4). We refer to these models as piecewise.

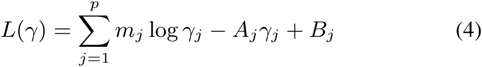

Here *A_j_* and *B_j_* are constants, for a given 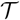, and 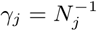. Taking partial derivatives we get 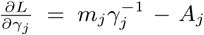 and observe that the MLE of 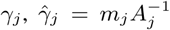. The second derivatives follow as: 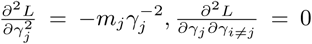. This leads to a diagonal Fisher information matrix 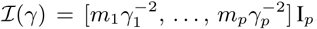, with I*_p_* as a *p* × *p* identity matrix. Using Eq. (1) we obtain the Fisher information in our original parametrisation as Eq. (5).

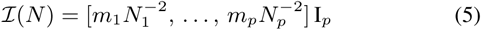

Several points become obvious. First, the achievable precision around 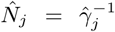 depends on the square of its unknown true value. This is a highly undesirable property, since it means estimation confidence will rapidly deteriorate as *N_j_* grows. Second, if our inference method directly worked in *γ*, instead of *N* (which is not uncommon for harmonic mean estimators [2]), then the region in which we achieve good *γ* precision is exactly that in which we obtain poor *N* confidence.

Third, the design variable *m_j_* only informs on one parameter of interest, *N_j_* or *γ_j_*. Good designs must therefore achieve *m_j_* ≥ 1 for all *j*. Failure to attain this will result in a singular Fisher information matrix and hence parameter non-identifiability [32], which can lead to issues like poor algorithmic convergence. This is particularly relevant for coalescent inference methods that feature pre-defined parameter grids of a size comparable to the tree size *n* [33].

Using either the *N* or *γ* parametrisation creates issues even when optimal design is employed. Consider the *N* parametrisation which has 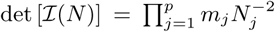. We let the constant 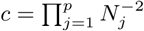. D-optimality is the solution to 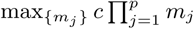 subject to 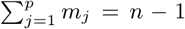. Our objective function is therefore 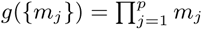 which is known to be Schur concave when all *m_j_* > 0. The optimal design is uniform and is the first equality in Eq. (6) below.

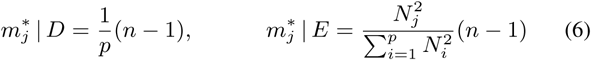

The E-optimal design solves: 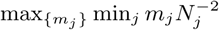. The objective function is now 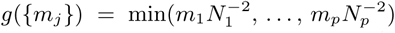 and is also Schur concave. The E-optimal solution satisfies 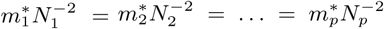 [31], and is the second equality in Eq. (6). This optimal design assigns more coalescent events to larger populations with a square penalty. The equivalent D and E-designs for inverse population size follow by simply replacing *N_j_* with *γ_j_* in Eq. (6) above.

Thus, in theory, D-optimal designs that consider *N* or *γ* could result in some parameters being very poorly estimated while E-optimal ones could allocate all of the coalescent events to a single parameter, increasing the possibility of non-identifiability. Additionally, for a given criterion, optimal *N_j_* and *γ_j_* designs can be contradictory. A robust design that is insensitive to both the parameter values and the choice of optimality criteria is therefore needed.

**Fig. 2:**
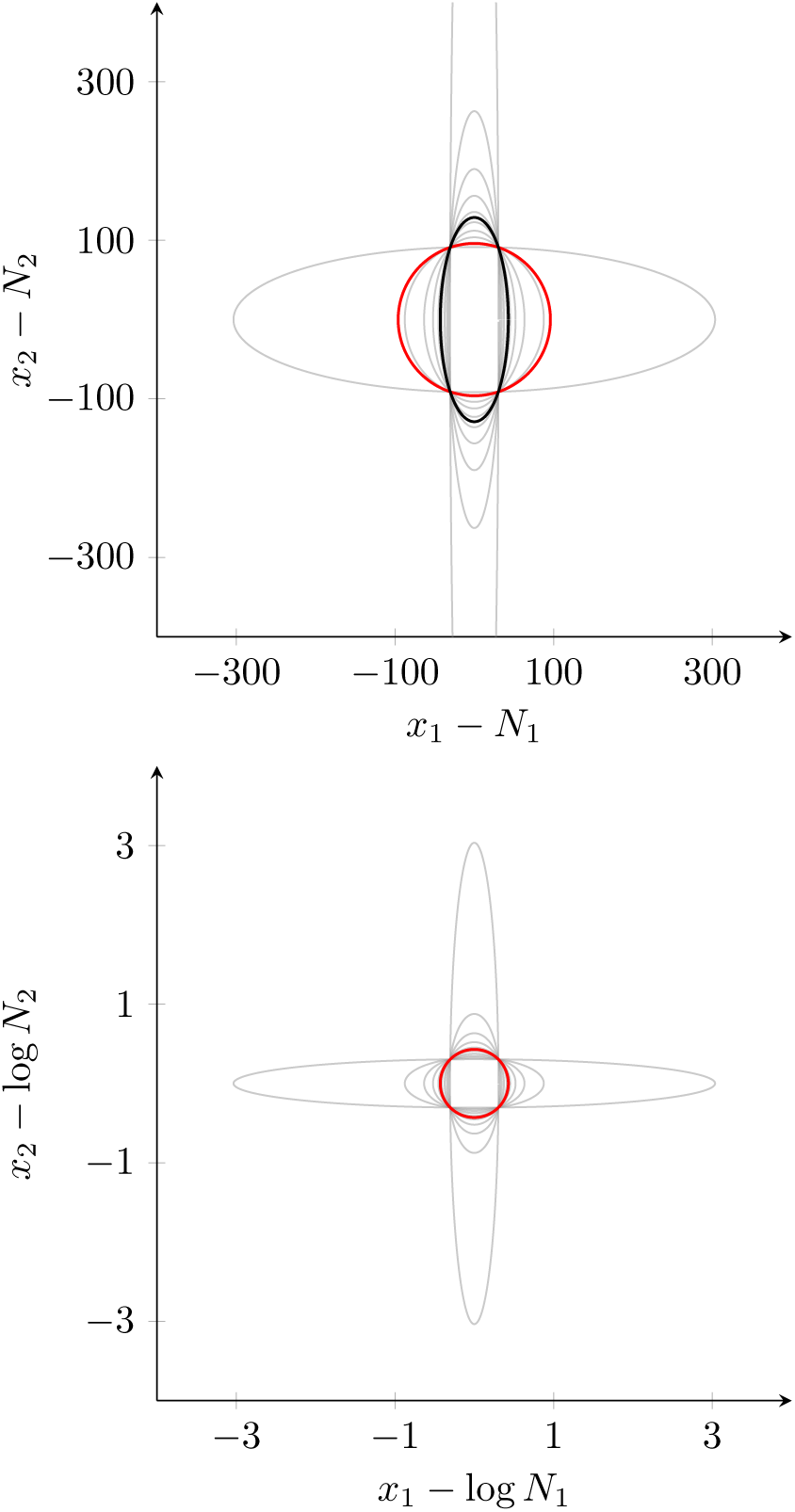
D and E-optimal designs for a two-parameter piecewise coalescent model. We provide asymptotic 99% confidence ellipses for a *p* = 2 skyline demographic design problem (see Fig. 1) with *n* − 1 = 100 = *m*_1_ + *m*_2_, *N*_1_ = 100 and *N*_2_ = 2*N*_1_. The ellipses depict the confidence region of the bivariate normal distribution that has covariance matrix equal to the inverse of the Fisher information. Each light grey ellipse indicates a different {*m*_1_, *m*_2_} distribution. D and E-optimal designs are in red and black respectively. The top panel shows the design space in absolute population size, *N_j_* with 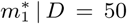 and 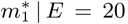. The bottom panel is in log population size, log *N_j_*, and leads to a symmetrical, robust design that has coincident D and E-optimal ellipses with 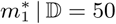.

This point is illustrated in the top panel of Fig. 2, which presents D and E-optimal confidence ellipsoids under *N*, for the model shown in Fig. 1. These ellipsoids, for some parameter vector *σ*, with diagonal Fisher information matrix 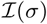, are given by 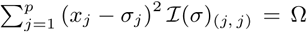. Here Ω controls the significance level according to a *χ*^2^ distribution (with *p* degrees of freedom) and *x_j_* is some coordinate on the *j*^th^ parameter axis [34]. Under *N* the D and E-optimal designs are notably different, and sensitive to the true values of *N*_1_ and *N*_2_.

#### Robust Coalescent Design

We define a robust experimental design as being (i) insensitive to the true (unknown) parameter values and (ii) minimising both the maximum and total uncertainty over the estimated parameters. The latter condition means that a robust design is also insensitive to choice of optimality criteria. We formulate our main results as the following two-point theorem. In subsequent sections we will apply this robust design to three popular coalescent models.

##### Theorem 1.

If the *p*-parameter vector *σ* admits a diagonal Fisher information matrix, 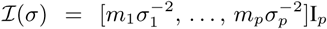, under an isoperimetric constraint 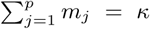, then any design that (i) works in the parametrisation [log *σ*_1_,…, log *σ_p_*] and (ii) achieves the distribution 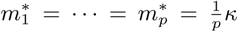 over this log *σ* space, is provably and uniquely robust.

Theorem 1 guarantees that inference is consistent and reliable across parameter space. We derive point (i), by maximising how distinguishable our parameters are within their space of possible values. ‘Distinguishability’ is a property that determines parameter identifiability and model complexity [35]. Let *ψ* be some parametrisation, in space Ψ, of a piecewise coalescent model. Then *h*(*ψ*) = *σ* defines a parameter transformation. Two vectors in Ψ, *ψ*_(1)_ and *ψ*_(2)_, are distinguishable if, given 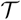, we can discriminate between them with some statistical confidence. Distinguishability is therefore intrinsically linked to the quality of inference. More detail on these information geometric concepts is given in [36] [35].

The number of distinguishable distributions in Ψ is described by the volume 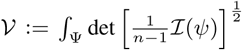. The *n* − 1 comes from the number of informative events in 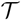. While 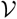 is invariant to the parametrisation choice *h* [35], different h functions control how the parameter space is discretised into distinguishable segments. For example, under *ψ* = *σ* poor distinguishability results when any *σ_j_* becomes large. We therefore pose the problem of finding an optimal bijective parameter transformation *h*(*ψ_j_*) = *σ_j_*, which maximises how distinguishable our distributions across parameter space are, or equivalently minimises the sensitivity or our estimates to the unknown true values of our parameters.

Applying Eq. (1), with 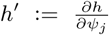, we get that 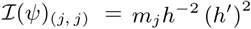. The orthogonality of the diagonal Fisher information matrix means that *ψ_j_* only depends on *σ_j_*. Using the properties of determinants, we can decompose the volume as 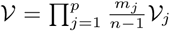. Since 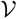 is constant for any parametrisation, our parameters are orthogonal and our transformation bijective, then 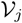 is also constant. If *σ_j_* ∈ [*σ_j_*_(1)_, *σ_j_*_(2)_], then *h*(*ψ_j_*_(1)_)= *σ_j_*_(1)_ and *h*(*ψ_j_*_(2)_)= *σ_j_*_(2)_. Using these endpoints and the invariance of 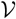 we obtain Eq. (7).

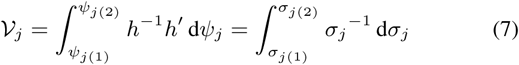

This equality defines the conserved property across parametrisations of coalescent models with likelihoods given in Eq. (4). We can maximise both the insensitivity of our parametrisation, *h*, to the unknown true parameters and our ability to distinguish between distributions across parameter space by forcing *h*^−1^*h*′ to be constant irrespective of *ψ_j_*. This is equivalent to solving a minimax problem. We choose a unit constant and evaluate Eq. (7) to obtain: *ψ_j_*_(2)_ − *ψ_j_*_(1)_ = log *σ_j_*_(2)_ − log *σ_j_*_(1)_. Due to the bijective nature of *h*, this implies that our (unique) optimal parametrisation is *ψ_j_* = log *σ_j_* and hence proves (i).

Point (ii) follows by solving optimal design problems under the log *σ* parametrisation. For consistency with Eq. (6), we set *σ* = *N*. This gives 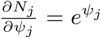 and results in the Fisher information matrix, 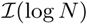, in Eq. (8).

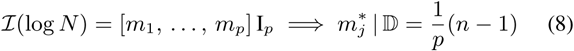

Let 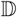 be an optimal design criterion, with event distribution 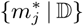. When 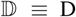, we maximise 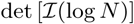 to obtain the uniform coalescent distribution in Eq. (8). The D-optimal design for *N*, *N*^−1^ and log *N* are therefore the same. However, we see interesting behaviour under other design criteria. When 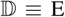, we maximise min 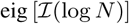 to again obtain Eq. (8). This is very different to analogous designs under *N* and *N*^−1^. While we do not assess further optimal design criteria here, several others also yield the design of Eq. (8).

Thus, under a log-parametrisation we see an important convergence of optimal experimental designs. This results in parameter confidence ellipsoids that are invariant to optimality criteria. This is shown in the bottom panel of Fig. 2 for a skyline model. This desirable design insensitivity emerges from the independence of 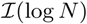 from *N*, for piecewise coalescent models, and proves (ii). We will now apply Theorem 1 to three different and commonly used coalescent models.

#### Skyline Demographic Models

Consider a coalescent process with deterministically time-varying population size, *N*(*t*), for *t* ≥ 0 that features sequences sampled at different times. As with the popular ‘skyline’ family of inference methods [2] [3] [4] [19], we assume that *N*(*t*) can be described by a piecewise-constant function with *p* ≥ 1 values so that 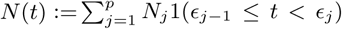 with *ϵ*_0_ = 0 and *ϵ_p_* = *∞*. *N_j_* is the constant population size of the *j*^th^ segment which is delimited by times [*ϵ_j_*_−1_, *ϵ_j_*). The indicator function 1(*a*) = 1 when *a* is true and 0 otherwise.

We start by assuming that this process has generated an observable coalescent tree, 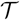, with *n* ≥ *n_s_* +1 tips, with *n_s_* ≥ 1 as the number of distinct sampling times. Each tree tip is a sample and the tuple (*s_k_*, *ϕ_k_*) defines a sampling protocol in which *ϕ_k_* tips are introduced at time *s_k_* with 1 ≤ *k* ≤ *n_s_* and 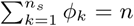. Since trees always start from the present then *s*_1_ = 0 and *ϕ*_1_ ≥ 2. In keeping with the literature, we assume that sampling times are independent of *N*(*t*) [4]. The choice of sampling times and the number of sequences obtained at each sampling time (i.e. the sampling protocol) is what the experimenter controls. Fig. 1 explains this notation for a *p* = 2 skyline demographic model.

The observed *n* tip tree has *n* − 1 coalescent events. We use *c_i_* to denote the time of the *i*^th^ such event with 1 ≤ *i* ≤ *n* − 1. We define *l*(*t*) as a piecewise-constant function that counts the number of lineages in 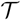 at *t* and let 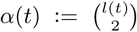. At the *k*^th^ sample time *l*(*t*) increases by *ϕ_k_* and at every *c_i_* it decreases by 1. The rate of producing coalescent events can then be defined as: 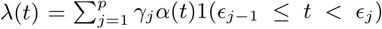 with 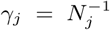 as the inverse population in segment *j*. We initially work in *γ* = [*γ*_1_,…, *γ_p_*], and then transform to *N* space.

The log-likelihood 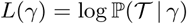 follows from Poisson process theory as [37] [5]: 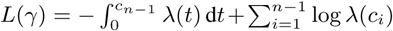. Splitting the integral across the *p* segments we get: 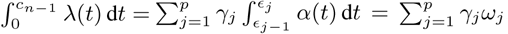. Here *ω_j_* is a constant for a given tree and it is independent of *γ*. Similarly, 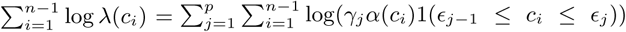. Expanding yields Eq. (9) with Γ*_j_* as a constant depending on *α*(*c_i_*) for all *i* falling in the *j*^th^ segment. The count of all the coalescent events within [*ϵ_j_*_−1_; *ϵ_j_*] is *m_j_*.

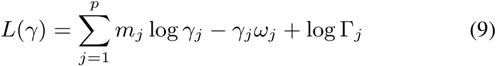

Eq. (9) is an alternate expression of the skyline log-likelihood given in [4], except that *N*(*t*) is not constrained to change only at coalescent event times. Importantly, sampling events do not contribute to the log-likelihood [4]. As a result we can focus on defining a desired coalescent distribution across the population size intervals, 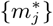. An optimal sampling protocol would then aim to achieve this coalescent distribution.

Since Eq. (9) is equivalent to Eq. (4), Theorem 1 applies. The relevant robust design is given by Eq. (8), and recommends inferring log *N* and sampling sequences in such a way that 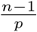 coalescent events fall in each [*ϵ_j_*_−1_, *ϵ_j_*) segment. Note that the number of lineages, *l*(*t*), the timing of the *m_j_* events within [*ϵ_j_*_−1_, *ϵ_j_*), and the wait between the last of these and *ϵ_j_* are all non-informative about population size. As an illustrative example, we solve a simple skyline model design problem in the Supporting Text. There we apply Theorem 1 to a square wave approximation of a cyclic population size function and find sampling protocols that achieve robust 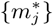 designs.

Lastly, we consider the impact of priors. More recent inference methods, such as the skyride [19] and skygrid [33] approaches, use smoothing priors that ease the sharpness of the inferred piecewise-constant population profile. While these priors embed extra (implicit) information about *N*, they do not alter the optimal design point, even for small *n*. This follows because the informativeness of a prior is unaffected by {*m_j_*} choices. The robust design therefore proceeds as above, independent of any contributions from the smoothing prior.

#### Structured Coalescent Models

Let 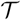 be an observed structured coalescent tree with *p* ≥ 1 demes that have been sampled through time (branches are labelled according to the deme in which they exist). Our experimental variables are the placement (both in time and in deme location) of the samples, and our goal is to define robust coalescent and migration design objectives. We set *T* as the number of intervals in 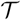, with each interval delimited by a pair of events, which can be sampling, migration or coalescent events. The *i*^th^ interval has length *u_i_* and 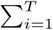 gives the time to the most recent common ancestor of 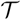. We use *l_ji_* to count the number of lineages in deme *j* during interval *i*. Lineage counts increase on sampling or immigration events, and decrement at coalescent or emigration events. We define the migration rate from deme *j* into *i* as *ζ_ji_*. *N_j_* and 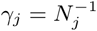 are the absolute and inverse population size in deme *j*.

Our initial *p*^2^ vector of parameters to be inferred is 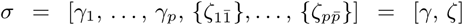, with 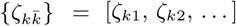 as the *p* − 1 sub-vector of all the migration rates from deme *k*. The log-likelihood 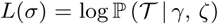 is then adapted from [21] and [38]. We decompose 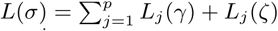 into coalescent and migration sums with *j*^th^ deme components given in Eq. (10) and Eq. (11). Here *m_j_* and *w_jk_* respectively count the total number of coalescent events in sub-population *j* and the sum of migrations from that deme into deme *k*, across all *T* time intervals. The factor 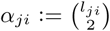 accounts for the contribution of the number of lineages to the coalescent rates. We constrain our tree to have a total of *n* − 1 coalescent events so that 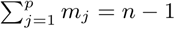.

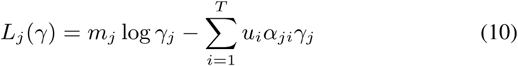

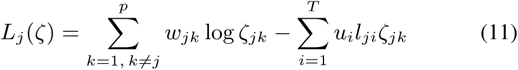

The log-likelihoods of both Eq. (10) and Eq. (11) are generalisations of Eq. (4) and lead to diagonal (orthogonal) Fisher information matrices like Eq. (5). This orthogonality results because migration events do not inform on population size and coalescent events tell us nothing about migrations. While migrations do change the number of lineages in a deme that can then coalesce, the lineage count component of the coalescent rate, *α_ji_*, does not influence the Fisher information. Importantly, since the Fisher information is independent of the sample times and locations, we can tune our sampling protocols to potentially achieve optimal design objectives.

Applying Theorem 1, we find that we should infer log population sizes and log migration rates from structured models. This removes the dependence on both the unknown population sizes and migration rates, and leads to a Fisher information of 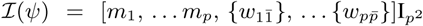 when 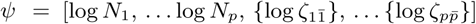. The robust design under this *ψ*, given in Eq. (12), involves distributing coalescent and migration events uniformly among the demes. Note that the migration rate distribution, 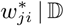, only holds if the total number of migration events are fixed, i.e. 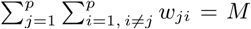, for some constant *M*.

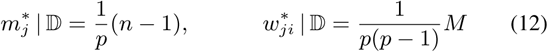

Two points are clear from Eq. (12). First, if all the migration rates are known, so that only population sizes are to be estimated then the structured model yields exactly the same robustness results as the skyline demographic model. Second, the migration rate design is the same at both the strong and weak migration limits of the structured model [39]. Thus, the true (unknown) migration rates do not affect their optimal design, provided log-migration rates are inferred.

If we generalise the population size function in each deme to be piecewise-constant in time, then we obtain a combination of the structured and skyline model design results. The robust design in this case maintains the log-population and migration recommendations but now requires that coalescent events are equally divided among both the demes and the piecewise-constant population segments.

#### Sequentially Markovian Coalescent Models

We now focus on coalescent models where recombination is applied along a genome, resulting in many hidden trees (multiple loci) [10]. Each tree typically consists of a small number of lineages. Popular inference methods in this field are based on an approximation to the coalescent with recombination called the sequentially Markovian coalescent (SMC) [40]. These methods typically handle SMC inference by constructing a hidden Markov model (HMM) over discretised coalescent time [10] [41] [11]. If we partition time into *p* segments: 0= *ϵ*_0_ < *ϵ*_1_ <… < *ϵ_p_* = *∞* then, when the HMM is in state *j*, the coalescent time is in [*ϵ*_j−1_, *ϵ_j_*) [11]. Recombinations lead to state changes in the HMM and the genomic sequence serves as the observed process of the HMM. Expectation-maximisation type algorithms are used to iteratively infer the HMM states from the genome [10] [41].

A central aspect of these techniques is the assumption that during each coalescent interval the population size is constant [12]. If the vector *N* = [*N*_1_,…, *N_p_*] denotes population size, then it is common to assign *N_j_* for the [*ϵ_j_*_−1_, *ϵ_j_*) interval [10]. This not only allows an easy transformation from the inferred HMM state sequence to estimates of *N* [13] but also controls the precision of SMC based inference. For example, if too few coalescent events fall within [*ϵ_j_*_−1_, *ϵ_j_*), then *N_j_* will generally be overestimated [11]. Thus, the choice of discretisation times (and hence population size change-points) is critical to SMC inference performance [12] [42].

Our experimental design problem therefore involves finding an optimal criterion for choosing these discretisation times. Currently, only heuristic strategies exist [11] [13] [12]. We define a vector of bins *β* = [*β*_1_,…, *β_p_*] such that *β_j_* = *ϵ_j_* − *ϵ_j_*_−1_ and assume we have *T* loci (and hence coalescent trees). In keeping with [10] and [41] we assume that each tree only leads to a single coalescent event, and hence we can neglect lineage counts. Since these counts merely rescale time (piecewise) linearly, we do not lose generality.

Let *m_ij_* be the number of coalescent events observed in bin *β_j_* from the locus so that 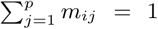. We further use 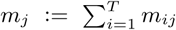 to count the total number of events from all loci falling in *β_j_*. As before, we constrain the total number of coalescent events so that 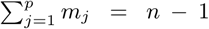. Using Poisson process theory we can write the log-likelihood of observing a set of coalescent event counts {*m_ij_*}, within our bins {*β_j_*} for the *i*^th^ locus as 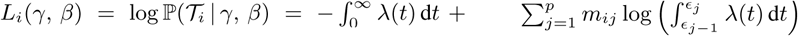 [37]. Here *λ*(*t*) is the coalescent rate at *t* so that 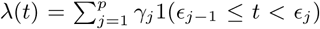 and 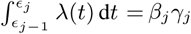 with 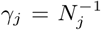. Using the independence of the *T* loci gives the complete log-likelihood of Eq. (13).

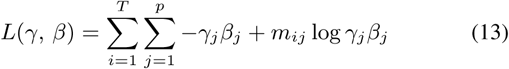

Eq. (13) is an alternative form of the log-likelihood given in [43], and describes a binned coalescent process that is analogous to the discrete one presented in [42]. Interestingly, Eq. (13) is a function of the product 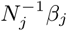 so that we cannot identify both the bins and the population size without extra information. This explains why choosing a time discretisation is seen to be as difficult as estimating population sizes [13].

Eq. (13) is analogous to Eq. (4), and so results in Fisher information matrices with square dependence on either *N_j_* or *β_j_* depending on what is known. Applying Theorem 1, we find that it is optimal to infer log-bin size (*ψ* = [log *β*_1_,…, log *β_p_*]), if population size history is known (this corresponds to discretisation results presented in [42]), or log-population size (*ψ* = [log *N*_1_,…, log *N_p_*]), if the bins are known. We generally assume the latter since bin end-points can often be set by the user [12]. Under either parametrisation, the provably robust design objective is to discretise time such that the resulting bins contain equal numbers of coalescent events.

## Discussion

Judicious experimental design can improve the ability of any inference method to extract useful information from observed data [44]. Despite these potential advantages, experimental design has received little attention in the coalescent inference literature [15]. We therefore defined and investigated robust designs for three important and popular coalescent models. Theorem 1, which summarises our main results, presents a clear and simple two-point robust design benchmark.

The first point recommends inferring the logarithm (and not the absolute value or inverse) of our parameters of interest. As this is usually effective population size, *N*, then log *N* is the uniquely robust parametrisation for piecewise coalescent estimation problems. While methods using log *N* do exist [12] [19], the stated reasons for doing so are centred around algorithmic convenience. Here we provide firm theoretical backing for using log *N* in coalescent inference.

The second point requires equalising the number of coalescent events informing on each parameter. This may initially appear obvious, as apportioning data evenly among the unknowns to be inferred seems wise. Indeed, [11] and [42], which focus on SMC models, state that time discretisations should aim to achieve uniform coalescent distributions. However, no proof for this statement is given. Here we not only provide theoretical support for uniform coalescent distributions, but also prove that they are only robust if the log-parameter stipulation is jointly satisfied.

Several unifying insights for piecewise coalescent models (i.e. those with likelihoods of form Eq. (4)) emerge as corollaries of our analysis. Because the precision with which we estimate a coalescent parameter only depends on the number of events informing on it, we can reinterpret all the designs considered here simply as different ways of allocating events to ‘pigeon-holes’. These pigeon-holes correspond to skyline intervals, structured coalescent demes and SMC time discretisation bins. This perspective reveals a straightforward rule for statistical identifiability: any piecewise coalescent model with at least one empty pigeon-hole is non-identifiable. This has specific ramifications. For example, it implies that we need at least one coalescent and migration event in each deme of the structured coalescent model to guarantee identifiability.

Knowing the boundaries or change-points of our pigeon-holes (e.g. the {*ϵ_j_*} for the SMC) is crucial for inference [42]. Throughout, we have assumed that these are indeed known. This is reasonable as it is generally not possible to jointly infer parameters and their change-points [11] [42]. Methods that do achieve this are usually data driven, iterative and case specific, allowing no general design insight [12] [45]. This raises the question about how to derive optimal design objectives when the change-points are unknown.

In the Supporting Text, we use Theorem 1 to compute robust change-point objectives. Interestingly, we show that it is wise to assign change-points according to the 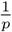 quantiles of the normalised lineages through time plot of the observed phylogeny. This results in a maximum spacings estimator (MSE) that makes the observed tree as uniformly informative as possible, relative to the pigeon-holes [46]. This means that if we wish to robustly infer *p* log-parameters from a tree containing *n* − 1 coalescent events, we should define our pigeon-holes such that they change every 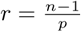 events.

Optimal skyline population profiles were examined in [3], with change-points selected on the basis of time. Our results suggest change-points should be based on coalescent event counts. If *r* =1, we recover the classic skyline plot [2] as the low information limit of this MSE strategy. Under our unifying corollary, grouping skyline intervals is analogous to aggregating demes in structured models, or combining bins in the SMC. Interestingly, this MSE strategy formally connects some popular SMC design choices. Specifically, [10] based its discretisation on a log spacing through time, while [41] used the quantiles of an exponential distribution. Our MSE result recommends using quantiles with logarithmic time bins.

Another unifying insight from Theorem 1 is that any parameter entering the coalescent log-likelihood in a functionally equivalent way to *γ_j_* in Eq. (4), should be inferred in log-space. This maximises distinguishability in model space, and means, for example, that it is best to work with log-migration rates for structured coalescent models. Using the log of the migration matrix is uncommon and could potentially improve current structured coalescent inference algorithms. Similarly, for the SMC, this insight implies that we must decide between absolute bin sizes for inferring log-populations and absolute population sizes for estimating log-bin widths.

Theorem 1 is also useful for finding cases where non-robust designs are inevitable. In the skyline demographic model, for example, a short interval during which population size is large would be difficult to estimate. Large *N* implies long coalescent times, making it unlikely that 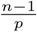 events can be forced to occur in such regions (see the square wave example in the Supporting Text). This hypothesis is corroborated by [13]. A similar effect occurs for SMC models if the bin size is small during a period of large population size [11]. For the structured model, the log-population size criteria is likely simpler to achieve than the log-migration rate one, since controlling *p*−1 stochastic migration event types per deme could be challenging, depending on how close the process is to the strong or weak migration limits [47] [48].

While we have provided robust coalescent design objectives, we have not defined what sampling or discretisation protocols can be used to achieve them. Existing analyses on this topic [14] [16] [47] [12] tend to examine a set of reasonable but ad-hoc protocols via extensive simulation. However, since no optimal design references exist, these works could only compare performance among their chosen protocols. Our analytical approach provides a general robust design theorem that can be used by future simulation studies for benchmarking.

## Acknowledgements

This work was supported by the European Research Council under the European Commission Seventh Framework Programme (FP7/2007–2013)/European Research Council grant agreement 614725-PATHPHYLODYN. The authors thank Louis Du Plesis and Chieh-Hsi Wu for their helpful comments.

## Supporting Text

### Robust Coalescent Change-point Designs

Consider the class of ‘piecewise’ coalescent models, which we define as having log-likelihoods analogous to Eq. (4) in the main text. This class includes the skyline demographic model, structured coalescent model and the SMC. We derived a robust design theorem (Theorem 1 of the main text) for inferring the parameters (e.g. effective population size) of these models. Theorem 1 suggested that experimental designs under piecewise coalescent models could be viewed as allocations of informative events (e.g. coalescent events) to ‘pigeon-holes’, which essentially encapsulate the different parameters that we wish to infer. These pigeon-holes, for example, are the piecewise-constant population size segments in the skyline demographic model, the demes of the structured coalescent model and the bins in the SMC. The boundaries or change-points of these pigeon-holes effectively control the complexity of our coalescent inference problem.

The analysis behind Theorem 1 presumed that we had knowledge of the pigeon-hole change-points. This corresponds to knowing the piecewise-constant segment times of the skyline model, the number of demes in the structured coalescent, and the bin sizes in the SMC. Such assumptions are reasonable, since simultaneously inferring both change-points and parameter values is an ill-conditioned problem. For example, if we do not know anything a-priori about either bin or population size, then it is impossible to derive optimal SMC time discretisations [11] [42]. Similar identifiability problems emerge when trying to simultaneously infer the change-points of piecewise-constant segments and their population sizes, or the number of demes and the population sizes and migration rates within each deme. In such cases iterative and data-driven computational methods can be employed [12] [45]. These methods will typically jointly optimise over these unknowns and produce sensible estimates, but their results will be case specific, allowing no general design insight to be derived.

While the general change-point inference problem is outside the scope of our work, we can provide some guidelines on how to robustly specify pigeon-hole change-points using the observed coalescent genealogy. We do this explicitly within the context of the SMC, but observe that the same results apply to all other piecewise coalescent models. It is known that if we condition on *n* − 1 events from an inhomogeneous Poisson process occurring in [0, *ϵ_p_*], with intensity *λ*(*t*), then the event times are independently and identically distributed according to density 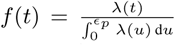 [37]. If we let *λ*(*t*) be our piecewise-constant SMC rate we find that 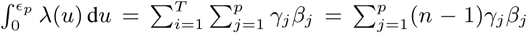, with 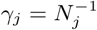 as the inverse population size over the region [*ϵ_j_*_−1_, *ϵ_j_*]. The pigeon-hole size or bin width is *β_j_* = *ϵ_j_* − *ϵ_j_*_−1_ with the *ϵ_j_* as the change-points, and *T* as the number of loci. Note that, for example, in the skyline demographic model, we would have a single locus and the *β_j_* would correspond to scaled interval times (see *ω_j_* in the derivation of the skyline demographic log-likelihood in the main text).

We can define the cumulative distribution function (CDF) at the pigeon-hole change-points as: 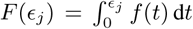 and denote the consecutive spacing of this CDF as Δ*_j_* = *F*(*ϵ_j_*) − *F*(*ϵ_j_*_−1_). Empirically, this CDF corresponds to the lineage through time plot (LTT) of the observed phylogeny, normalised by its total number of coalescent events. Solving for Δ*_j_* using the piecewise-constant coalescent rate gives the left part of Eq. (14). This expression is precisely the same for the skyline and structured models. If we substitute the MLE for either *β_j_* or *γ_j_* (depending on what is known) then we derive 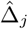. Applying the 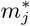 design from Theorem 1 produces the rest of Eq. (14).

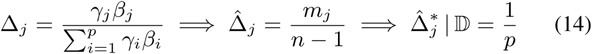

The robust coalescent interval spacing, 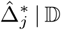, is therefore fixed by the number of pigeon-holes (and hence parameters). This has two important ramifications. First, as quantiles are defined as inverse cumulative distribution values, it means that the optimal choice of pigeon-holes is such that their boundaries are the 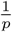 quantiles of the normalised LTT. Robust coalescent experimental design therefore recommends assigning a new pigeon-hole after every 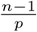 events of the LTT. This quantile design clearly suggests that the largest admissible number of change-points occurs when *p* = *n* − 1. This limit, for skyline demographic inference problems, corresponds to the formulation of the classic skyline plot [2].

Second, since the spacing at the MLE is constant, robustness is achieved by the maximum spacings estimate (MSE) [49] [46]. For a given set of observations, drawn from the CDF of a parameter *θ*, the MSE is the estimate of *θ* that maximises the geometric mean of the spacing of the CDF, evaluated at each observed random sample. Our results suggest that if we view the pigeon-hole change-points as binned draws from *f*(*t*) then, given a robust design, the MSE of *θ* results in optimal spacing. Here *θ* is the effective coalescent rate with density *f*(*t*). It is not difficult to prove that robust designs for the skyline demographic and structured models also imply equivalent 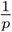 MSEs. Under MSE designs, the observed tree, from the perspective of the pigeon-holes, will appear as uniformly informative as possible.

### Simulation Study: Square Wave Populations

Here we show how to apply Theorem 1 to a simple skyline demographic coalescent model. Let *N*(*t*), define a square wave population size function with period *T*, with time *t* into the past. *N*(*t*) models the harmonic mean [2] of the fluctuating number of infected individuals across time in a seasonal epidemic. *N*_1_ recurs on odd half-periods and *N*_2_ on even ones 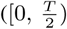 is the first (odd) half-period). Given *n* total samples (*n* − 1 coalescent events) we want to optimally infer *N*(*t*). Fig 1 of the main text illustrates the experimental set-up and notation for a similar design problem. Panel (a) of Fig. 3 shows a typical *N*(*t*) with its half-period numbers.

**Fig. 3:**
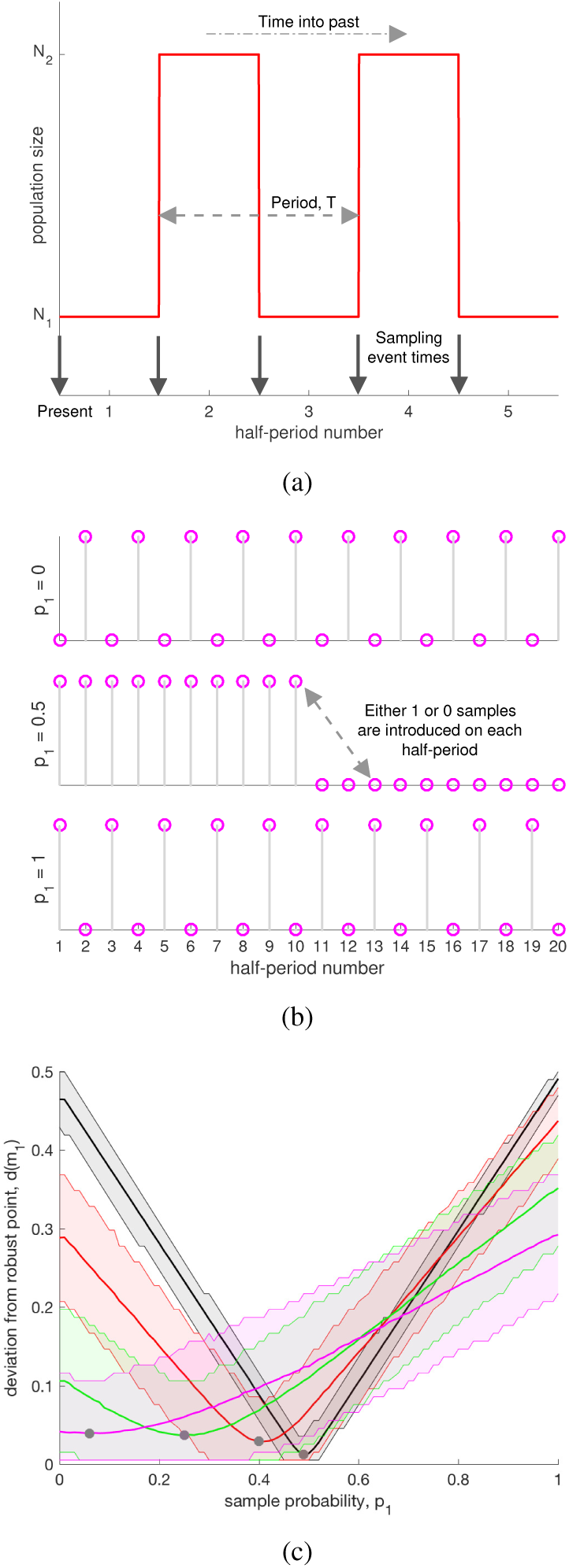
Deterministic sampling protocols for a skyline coalescent model. We apply a deterministic sampling strategy with *ϕ_i_* = 1 or 0 to a skyline demographic model with a population that fluctuates between *N*_1_ and *N*_2_ = 2*N*_1_ across time. This fluctuation is described by a square wave with period *T*, and is shown in panel (a) for 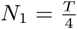 and 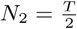. The arrows in this sub-plot indicate the points at which we can introduce a sample. Panel (b) shows how *n* = 10 samples are allocated at these arrow points for three different *p*_1_ protocols (*p*_1_ controls the fraction of the *n* available samples that are placed in *N*_1_ half-periods). We observe how the absolute difference, *d*(*m*_1_), between the Fisher information and the uniquely robust design changes with *p*_1_ in panel (c), for *n* = 100. The black, red, green and magenta curves are for 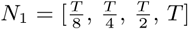 respectively. Each curve gives the mean of *d*(*m*_1_) across 5000 repeated runs (solid line) and the 95% confidence interval around that mean. As *N*_1_ decreases relative to *T*, *d*(*m*_1_) becomes more symmetrical and maximal performance (defined as min *d*(*m*_1_)) improves (gets closer to 0 and has sharper confidence). The uniquely robust sampling protocol in each *N*_1_ case, is visualised with a grey, filled circle. See the supporting text for further interpretations of these results.

The precision with which *N*_1_ and *N*_2_ are estimated is an increasing function of the number of coalescent events falling within their half periods. Let 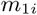 be the number of events in the *i*^th^ recurrence of *N*_1_ and 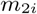 be the equivalent for *N*_2_. Theorem 1 stipulates that robust sampling schemes will distribute 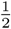 of all coalescent events to *N*_1_ half-periods (Eq. (15)). Thus, if *m*_1_ is the observed count of coalescent events falling within *N*_1_ half-periods, then the performance of any sampling scheme can be measured by the size of the scalar 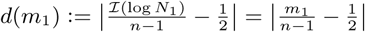. Note that *d*(*m*_1_) increases as the Fisher information becomes more skewed (higher 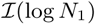 means lower 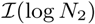), and 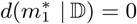.

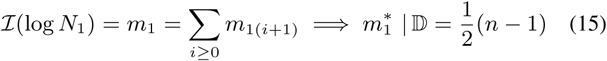

If we define, *p*_1_, as the probability that a sampled tip is introduced in an *N*_1_ interval then a robust sampling strategy achieves 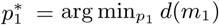. We assume *p*_1_ is constant with time. Thus, we focus on the mapping *p*_1_ → *d*(*m*_1_) with *p*_2_ =1 − *p*_1_. A sampling protocol involves the tuple (*s_k_*, *ϕ_k_*) with *s_k_* as the time of the *k*^th^ sampling event at which *ϕ_k_* lineages are introduced. Since coalescent events are always delayed in time relative to the point in time at which samples are placed, we will always introduce our *ϕ_k_* samples all at once, and only at the change-points so that 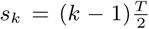 (the arrows in panel (a) of Fig. 3). This procedure maximises the probability that samples will coalesce within the half-period in which they are introduced.

We examine a range of deterministic sampling strategies in order to explore how *p*_1_ controls *d*(*m*_1_). For a given *p*_1_, we set the number of samples introduced in *N*_1_ and *N*_2_ half-periods as fractions *f*_1_ = round [*p*_1_(*n* − 1)] and *f*_2_ = *n* − 1 − *f*_1_. Here round indicates the nearest integer. Note that 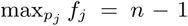 as we assume that there is always an initial sample to allow the first coalescent event. We allocate the *f*_1_ and *f*_2_ samples uniformly relative to *N*_1_ and *N*_2_ half-periods respectively, so that *ϕ_i_* = *a* or 0 depending on whether samples are introduced or not. Here *p*_1_ = 0 means we have placed all *n* samples on *N*_2_ half-periods while *p*_1_ = 1 means that they are all on *N*_1_ ones. Intermediate *p*_1_ values compromise between these two extremes. We illustrate these sampling strategies for *a* = 1 and *n* = 10, relative to the half-periods of *N*(*t*), in panel (b) of Fig. 3.

Panel (c) of Fig. 3 shows the sampling protocol performance under *a* = 1 schemes at different *N*_1_ values (scaled against *T*), with *N*_2_ = 2*N*_1_. We find that as *N*_1_ becomes smaller relative to *T*, the optimal protocol 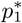 gets closer to 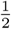. This makes sense since here population changes are slow relative to the coalescent times, so that we have the greatest chance of any sample falling within the half-period in which it was introduced. As *N*_1_ increases, coalescent times lengthen and we get samples falling outside this original half-period. This leads to a weaker, less discernible minimum with larger uncertainty (we cannot estimate fluctuations in population that are fast compared to our rate of producing coalescent events [48]). The optimal strategy here is 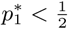 (if we made 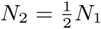 we would get curves skewed in the opposite direction so that 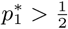). Robust sampling therefore favours placing more samples in periods of time with larger population size. This has an interesting implication for structured coalescent models with known, symmetrical migration rates. In this case the demes are directly analogous to the *N_j_* segments and robust sampling would recommend allocating sample numbers in proportion to the deme population sizes.

